# Widespread Alterations in Translation Elongation in the Brain of Juvenile *Fmr1* Knock-Out Mice

**DOI:** 10.1101/319368

**Authors:** Sohani Das Sharma, Jordan B. Metz, Hongyu Li, Benjamin D. Hobson, Nicholas Hornstein, David Sulzer, Guomei Tang, Peter A. Sims

## Abstract

FMRP is a polysome-associated RNA-binding protein encoded by *Fmr1* that is lost in Fragile X syndrome. Increasing evidence suggests that FMRP regulates both translation initiation and elongation, but the gene-specificity of these effects is unclear. To elucidate the impact of *Fmr1* loss on translation, we used ribosome profiling for genome-wide measurements of ribosomal occupancy and positioning in the cortex of 24 day-old *Fmr1* knock-out mice. We found a remarkably coherent reduction in ribosome footprint abundance per mRNA for previously identified, high-affinity mRNA binding partners of FMRP, and an increase for terminal oligo-pyrimidine (TOP) motif-containing genes canonically controlled by mTOR-4EBP-eIF4E signaling. Amino acid motif- and gene-level analyses both showed a widespread reduction of translational pausing in *Fmr1* knock-out mice. Our findings are consistent with a model of FMRP-mediated regulation of both translation initiation through eIF4E and elongation that is disrupted in Fragile X syndrome.

## Introduction

Fragile X syndrome (FXS) is a highly penetrant, heritable form of intellectual disability that is associated with autism. The most common cause of FXS is epigenetic silencing of the *FMR1* gene that encodes the fragile X mental retardation protein (FMRP). FMRP is an RNA binding protein that regulates both translation initiation and elongation (Darnell et al., 2011; Khandjian, 1999; Napoli et al., 2008; Stefani et al., 2004). Translation of the majority of cellular mRNAs begins with recognition of the of the 5’ cap structure m^7^G(5’)ppp(5’)N by eukaryotic initiation factor 4E (eIF4E). FMRP has been shown to repress translation initiation by interacting with cytoplasmic FMRP-interacting protein 1 (CYFIP1) (Napoli et al., 2008), an eIF4E binding protein which competes with eIF4G for interaction with eIF4E and prevents formation of the initiation complex (Richter and Sonenberg, 2005).

FMRP co-sediments with actively translating ribosomes and polyribosomes in gradient fractionation assays (Feng et al., 1997; Khandjian et al., 1996; Stefani et al., 2004). Recently, a genome-wide analysis of RNA-FMRP interactions was undertaken in the murine brain with high throughput cross-linking immunoprecipitation (HITS-CLIP) (Darnell et al., 2011). In this study, FMRP was found to bind primarily to protein-coding sequences (CDS) of mRNAs, and no specific binding motif was identified. The highest-affinity mRNA binding partners were enriched in postsynaptic and autism-related genes, including components of the mGluR5 metabotropic glutamate receptor complex and downstream PI3K signaling regulator PIKE, both of which are dysregulated in Fragile X Syndrome (Bear et al., 2004; Gross et al., 2015). *In vitro* puromycin run-off experiments on a set of nine high-affinity binding partners showed extensive, FMRP-dependent ribosomal stalling compared to genes with lower HITS-CLIP signal. Furthermore, the *in vitro* ribosome translocation rate was shown to be significantly higher in brain lysates of *Fmr1* knock-out (*Fmr1*-KO) mice than wild-type mice (Udagawa et al., 2013). Studies have also shown elevated rates of protein synthesis in brains of *Fmr1*-KO mice (Qin et al., 2005) and increased protein expression of many FMRP high-affinity mRNA binding partners (Tang et al., 2015). Taken together, these studies suggest that FMRP represses protein synthesis at the level of translation elongation by acting as a ribosomal brake.

Despite this progress, important questions remain regarding the nature of translational regulation by FMRP in the brain. While Darnell and colleagues have identified high-affinity binding partners, it is unclear whether there is a relationship between FMRP affinity and translational repression. Furthermore, it remains unknown whether FMRP represses translation in the brain through a dominant mechanism or whether both initiation and elongation are significantly affected. Ribosome profiling enables genome-wide measurement of ribosome density on mRNAs with single-nucleotide resolution, allowing simultaneous analysis of the overall ribosome density on each gene and ribosomal stalling. In this study, we conducted ribosome profiling and RNA-Seq in wild type and *Fmr1*-KO mice to obtain an unbiased, high-resolution assessment of the impact of *Fmr1* loss on protein synthesis in the brain.

## Results

### Translational landscape of Fmr1 knock-out mice

To determine the effect of *Fmr1* loss on translation, we conducted ribosome profiling and RNA-Seq on the frontal cortex of *Fmr1*-KO and wild-type male mice at postnatal day 24 (P24). Genome-wide ribosome footprint (RF) and RNA-Seq data were highly reproducible across biological replicates with genotype as the principal source of variation (**Figure 1A-B**). As expected, alterations in RF abundance and RNA expression were generally correlated (**Figure 1C, Supplementary Tables 1-2**), and a handful of genes exhibited particularly large differences in RF abundance between genotypes. For example, we found immediate early genes, including *Arc*, *Fos*, and *Egr2* to have significantly elevated RF abundance in *Fmr1*-KO mice, with much smaller alterations at the RNA level. To characterize the effects of *Fmr1* loss more broadly, we conducted differential RF abundance and RNA expression analyses. We used gene set enrichment analysis (GSEA) to assess differentially translated and expressed gene ontologies (GOs). Interestingly, while GOs associated with protein synthesis had higher RF abundance in *Fmr1*-KO mice compared to wildtype, translation-associated GOs exhibited lower expression at the RNA level in the *Fmr1*-KO mice (**Figure 1D-E**). In addition, GOs associated with neuronal projection development, morphology, and extracellular matrix have lower RF abundance in *Fmr1*-KO mice.

**Figure 1:**
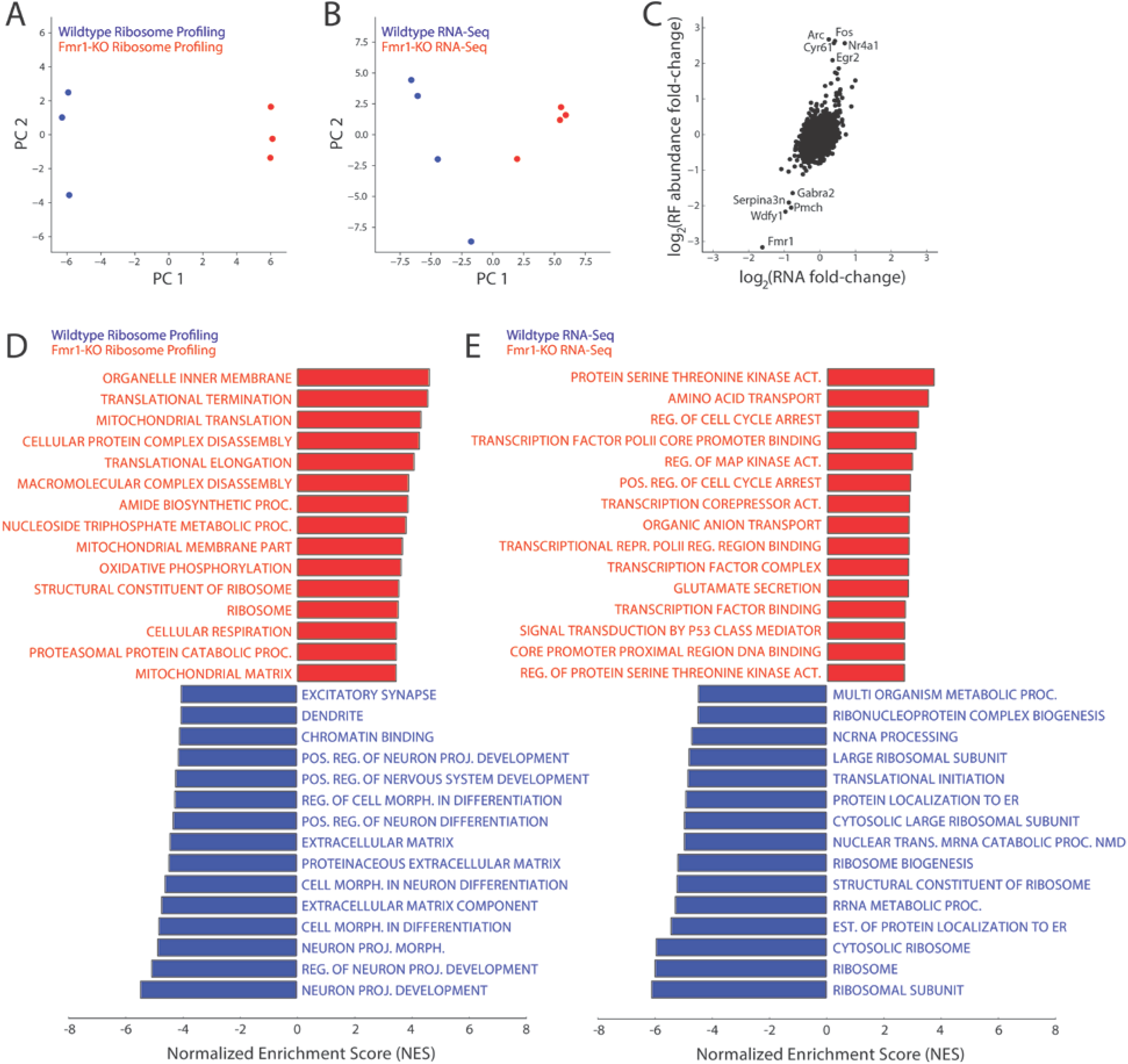
Principal component analysis (PCA) of both ribosome profiling (A) and RNA sequencing (B) libraries from *Fmr1*-KO and wild-type mice. Samples are segregated by genotype in principal component 1 (PC1), the axis representing the major source of variation in the data, in both plots. (C) Comparison of differential ribosome footprint abundance against differential RNA expression levels between genotypes at the level of individual genes. Though ribosome footprint abundance displays a greater range of changes than RNA expression level, these measurements are highly correlated. *Fmr1*, knocked out at the transcript level (by deletion of one exon), shows decreased RNA expression and ribosome density as expected, while the immediate early genes Fos, Arc, and Egr2 show increased ribosome density and RNA expression. (D,E) Enrichment scores of the top 15 gene ontologies (GOs) enriched in the wild-type or *Fmr1*-KO brain, determined by GSEA on genes ranked by their fold-changes in ribosome footprint abundance (D) or RNA expression (E) as presented in (C). Genes related to protein synthesis are enriched in ribosome density in *Fmr1*-KO mice compared to wildtype but depleted in RNA expression level (and therefore enriched for in wild-type vs *Fmr1*-KO mice), while ontologies related to neuronal development and morphology show decreased ribosome density in *Fmr1*-KO mice.

To identify genes with significant alterations in ribosome footprint abundance per mRNA (RFApm), calculated as the ratio of RF abundance and RNA expression, we used the generalized linear model (GLM) implemented in RiboDiff for joint statistical analysis of the ribosome profiling and RNA-Seq data (**Supplementary Table 3**).This metric approximates the number ribosomes bound per mRNA and is commonly referred to as “translation efficiency” (Ingolia et al., 2009). However, RFApm depends on complex relationships between the rates of translation initiation, elongation, and termination that complicate its interpretation (Arava et al., 2005). GSEA revealed that genes involved in protein synthesis have elevated RFApm in *Fmr1*-KO mice with concomitant reductions in genes associated with extracellular matrix and neuronal function, differentiation, and projection (**Figure 2A**). Translation initiation for effectors of protein synthesis such as ribosomal proteins and translation factors is regulated by mTOR signaling through a cis-regulatory element known as the 5’-terminal oligopyrimidine (5’TOP) motif found in the corresponding mRNAs (Hsieh et al., 2012; Thoreen et al., 2012). This regulation is mediated by 4E-BPs, which, in their dephosphorylated state, sequester the initiation factor eIF4E (Thoreen et al., 2012). FMRP can repress translation via an inhibitory FMRP-CYFIP1-eIF4E complex (Napoli et al., 2008; Santini et al., 2017) and *Fmr1*-KO mice exhibit increased eIF4E-dependent translation (Sharma et al., 2010). Therefore, we expected that the 5’TOP motif-containing mRNAs would exhibit increased RFApm in *Fmr1*-KO mice. Indeed, **Figure 2B** shows that the 5’TOP transcripts exhibited significantly higher RFApm in *Fmr1*-KO mice (p<0.00001, GSEA), consistent with a previously characterized mechanism through which FMRP modulates translation initiation.

**Figure 2:**
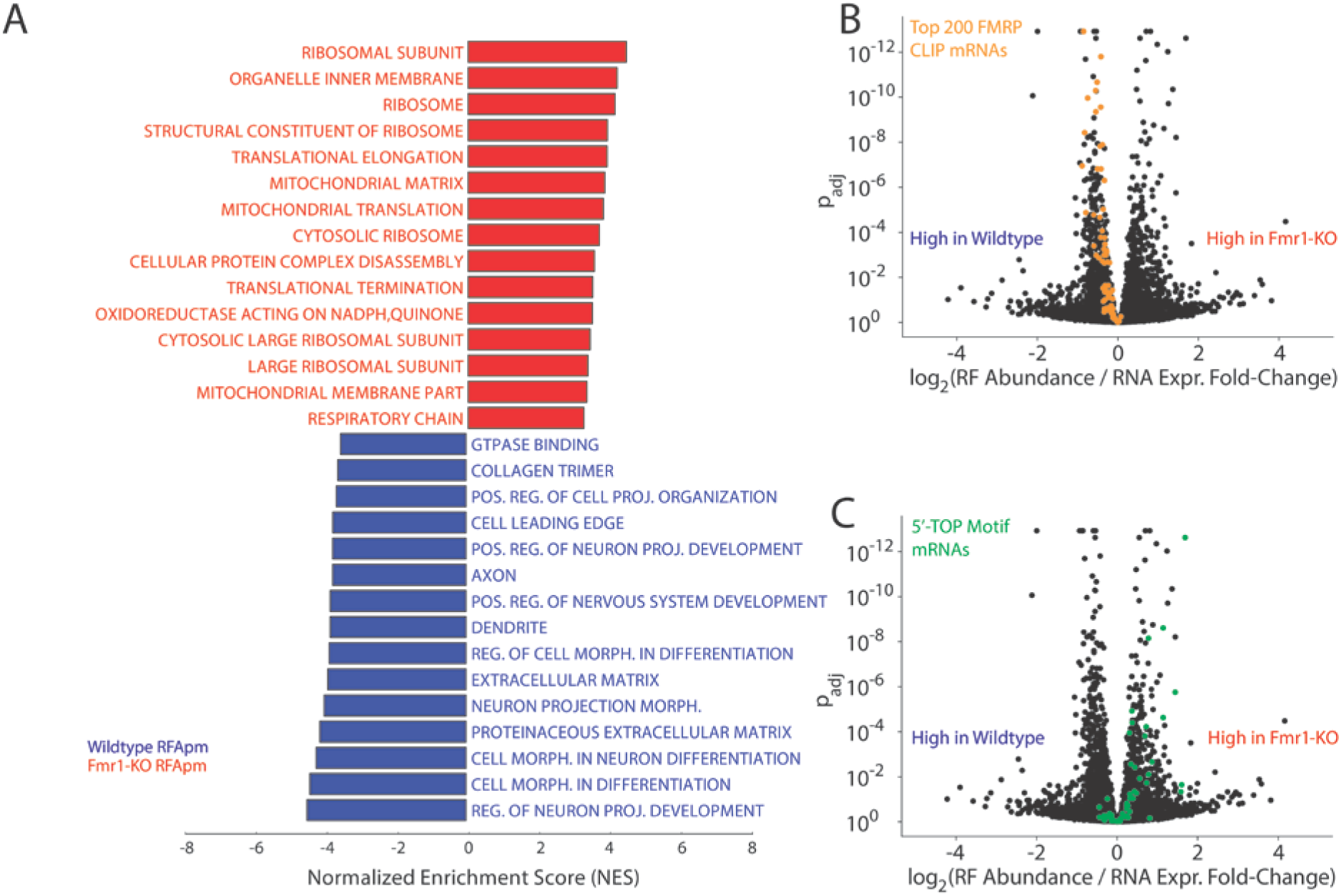
(A) GSEA performed on genes ranked by their differential ribosome footprint abundance per mRNA (RFApm) between genotypes reveals increased RFApm of genes related to protein synthesis (ribosome, translation elongation, mitochondrial translation) in *Fmr1-*KO mice with decreased RFApm of genes involved in neuronal projection development and morphology. (B) and (C) are volcano plots comparing the observed effect size of log-fold change in RFApm with adjusted p-values for all detected genes. (B) demonstrates a uniform, modest reduction in RFApm (p<0.00001, GSEA) across the top 200 highest-affinity binding partners for FMRP determined by HITS-CLIP in *Fmr1*-KO mice (orange), while (C) shows a trend towards increased RFApm in the 5’-terminal oligopyrimidine motif-containing (5’-TOP) genes (p<0.00001, GSEA), the canonical targets of mTOR (green). (B) and (C) together demonstrate the concerted dysregulation of distinct gene sets in opposite directions associated with FMRP loss.

As described above, earlier work showed that FMRP binds to mRNAs that encode proteins associated with synaptic activity and other neuronal functions. The GSEA in **Figure 2A** suggests a reduction in RFApm for genes with similar functions. Indeed, **Figure 2C** shows that the top 200 highest-affinity FMRP binding partners exhibit significantly reduced RFApm (p<0.00001, GSEA). This coherent reduction in apparent translation efficiency is surprising, because many of these genes have been shown to be over-expressed at the protein level in the brains of *Fmr1*-KO mice (Hou et al., 2006; Schutt et al., 2009; Zalfa et al., 2003; Zhang et al., 2001). One possibility is that protein synthesis from these mRNAs is controlled at the level of translation elongation. For example, a decrease in RFApm could result from a reduction in ribosomal stalling rather than in initiation efficiency (Ingolia et al., 2009).

### Alterations in translation elongation in Fmr1 knock-out mice

Given the previous evidence of FMRP-dependent ribosomal pausing (Darnell et al., 2011) and the results described above, we next quantified ribosomal pausing using the ribosome profiling data. Specifically, we calculated the ribosome pause score at the level of encoded amino acid sequences, averaging scores across all occurrences of codons corresponding to a given amino acid residue. This metric allows the determination of pause activity due to encoded peptide sequence. **Figure 3** compares the distributions of pause scores across mono-, di-, and tri-amino acids between *Fmr1*-KO and wild-type ribosome occupancy profiles. With few exceptions, sequences exhibited a lower mean pause score in *Fmr1*-KO than in wild-type profiles, demonstrated by a downward shift away from the main diagonal in **Figure 3A-C**. This shift indicates a global relief of pausing associated with *Fmr1* loss that is inconsistent with an effect on a limited set of specific binding partners.

**Figure 3:**
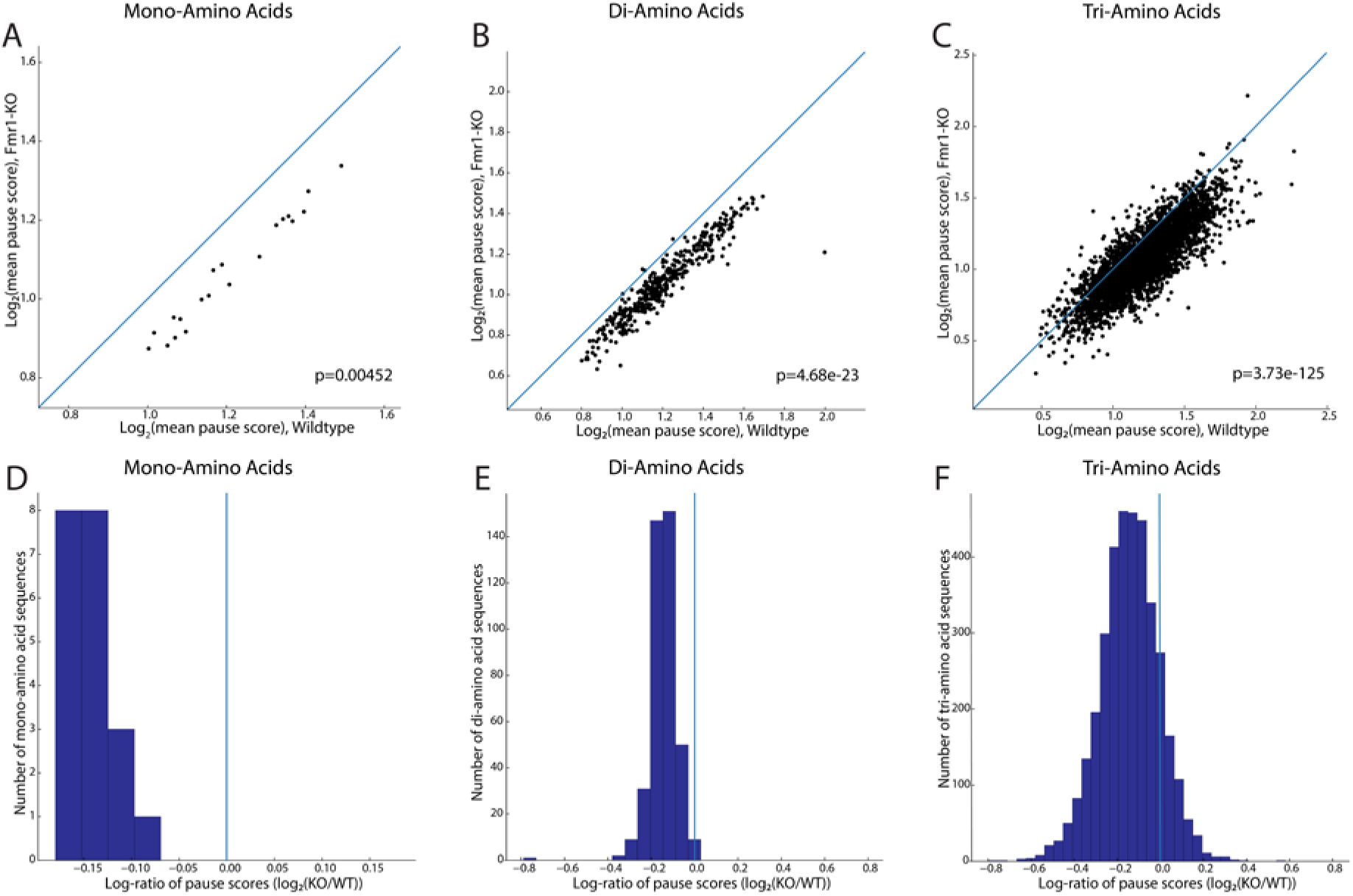
Log-log plots of mean pause scores calculated for single amino acid (A), di-(B), and tri-amino acid sequences (C) in *Fmr1*-KO and wild-type mice, with accompanying p-value for the significance of the difference in these two distributions (Mann-Whitney U-test). In each plot, the main diagonal is plotted as a blue line representing equal pausing in either genotype, highlighting the downward shift of the mass of individual sequences’ scores and decrease in pause score in *Fmr1*-KO mice. This shift is visualized differently in (D-F), histograms of the log-ratios of mean pause scores for every mono-(D), di-(E), and tri-amino acid motif (F). The downward/rightward shift in (A-C) translates to a leftward shift away from the blue vertical line at x=0, showing decreased pausing for the majority of encoded amino acid motifs in *Fmr1*-KO vs wild-type mice.

While codon-level analysis suggests that alterations in translation elongation are widespread, we further validated these changes directly at the gene-level. Gene-level analysis of translational pausing is complicated by the large dynamic range in gene expression, which results in a broad coverage distribution for ribosome profiling across genes. For example, consider two genes with similar translational pausing behavior where one gene is lowly expressed, resulting in a low-coverage ribosome profile. A naïve analysis might conclude that this lowly expressed gene has more translational pausing – an artifact of sparse coverage. At low coverage, it is challenging to differentiate noise (which scales inversely with coverage due to counting statistics) from real translational pausing. To address this issue, we developed an analytical method for gene-level analysis of translational pausing that explicitly models the dependence of noise in ribosome profiles on coverage.

**Figure 4A-C** shows the dependence of the noise (expressed as coefficient of variation or *CV*) in the ribosome profile along the CDS of each gene on coverage (expressed as ribosome footprint reads per codon). As expected, the *CV* decreases with increasing coverage regardless of genotype (**Figure 4A-B, Supplementary Figure 1**). We fit the following two-parameter model to the data to accommodate a variety of statistical behaviors for counting noise:

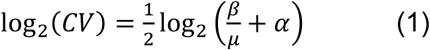

where *CV* is the coefficient of variation in the ribosome profile of a given gene, *μ* is mean coverage (ribosome footprint reads per codon), and *α* and *β* are fitting parameters. Importantly, when *α* = 0 and *β* = 1, Equation 1 results from a Poisson distribution whereas *α* > 0 and *β* = 1 indicates a negative binomial distribution. **Figure 4C** shows the fits for all wild type (n=3) and *Fmr1*-KO (n=3) ribosome profiling data sets. While biological replicates of each genotype are highly reproducible, there is a clear difference between genotypes with the *Fmr1*-KO mice exhibiting markedly lower *CV* at higher coverage. Over-dispersion is widely appreciated for RNA counting data derived from high-throughput sequencing, and as expected, *α* > 0 for all data sets. For highly translated genes, where coverage is drawn from an over-dispersed distribution, *CV* converges to *α*^1/2^. However, there is a strong genotype effect on *α*, (2.61±0.02 for wildtype and 1.720±0.002 for *Fmr1*-KO, p = 0.0001). Taken together, these results indicate that the ribosome profiles of genes in *Fmr1*-KO brains display less variability in coverage along the CDS than in the wildtype. These findings are consistent with the codon-level analysis described above, reflecting a global reduction in translational stalling in *Fmr1*-KO mice.

**Figure 4:**
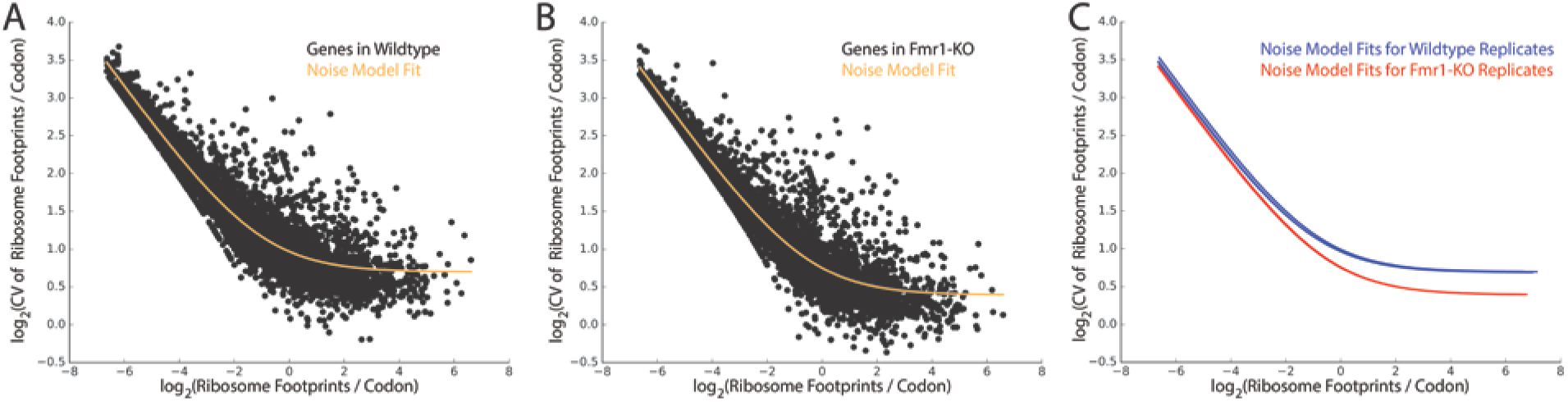
For gene-level analysis of ribosomal pausing, (A) and (B) plot the relationship between noise and coverage for a single wild-type and *Fmr1*-KO replicate, respectively. In these plots, coverage is the log-mean number of ribosome footprints aligned per codon of a given transcript, noise is represented by log-coefficient of variation, or standard deviation in the number of ribosome footprints per codon divided by mean, and the relationship of these values across each gene is summarized by regression to the two-parameter model in Equation 1, plotted in orange. (C) regression curves for each replicate on the same axes (n=3 for both genotypes), showing both the uniformity of this relationship across biological replicates, as well as the global downward shift in coefficient-of-variation of *Fmr1*-KO replicates relative to wild-type. This shift, representing a lower degree of coverage variation along the gene body, indicates a widespread reduction in pausing across transcripts.

The analysis in **Figures 3-4** suggests that loss of *Fmr1* results in widespread alterations in translation elongation. Although many of the high-affinity FMRP binding partners and 5’TOP motif-containing mRNAs display decreased and increased RFApm, respectively, nearly all of these genes exhibit reduced pausing in *Fmr1*-KO mice (**Supplementary Figure 2)**. **Figure 5** shows specific examples of this among representative genes from a few different categories. Importantly, there is not a large difference in coverage between wildtype and *Fmr1*-KO for any of these genes. **Figure 5A-B** show the P-site ribosome profiles for all three wildtype and *Fmr1*-KO mice for two genes with a significant reduction in RFApm. *Syn1* is a high-affinity FMRP binding partner (**Figure 5A**), while *Map1b* (**Figure 5B**) is not. There are two particularly notable features of these data. First, the *Fmr1*-KO profiles display a clear reduction in the large, reproducible pauses manifested as “spikes” in the wild type profiles. Second, as shown in the rightmost panel of **Figure 5A-B**, there is a reproducible, overall reduction in the *CV* along the gene body that is not explained simply by the reduction in large pauses. **Figure 5C-D** shows the same analysis for two genes with a significant increase in RFApm in *Fmr1*-KO mice. *Rpl4* is a TOP-motif gene, which is enriched among genes with an apparent increase in translation efficiency as described above, and the other (*Ndel1)* is not. For all four genes in **Figure 5**, we detect stereotyped pauses in the wild type that are substantially ablated in the knock-out. We also find a reproducible reduction in cumulative *CV* along the gene body, suggesting a smoother overall translocation process for the ribosome in the brain of *Fmr1*-KO mice.

**Figure 5:**
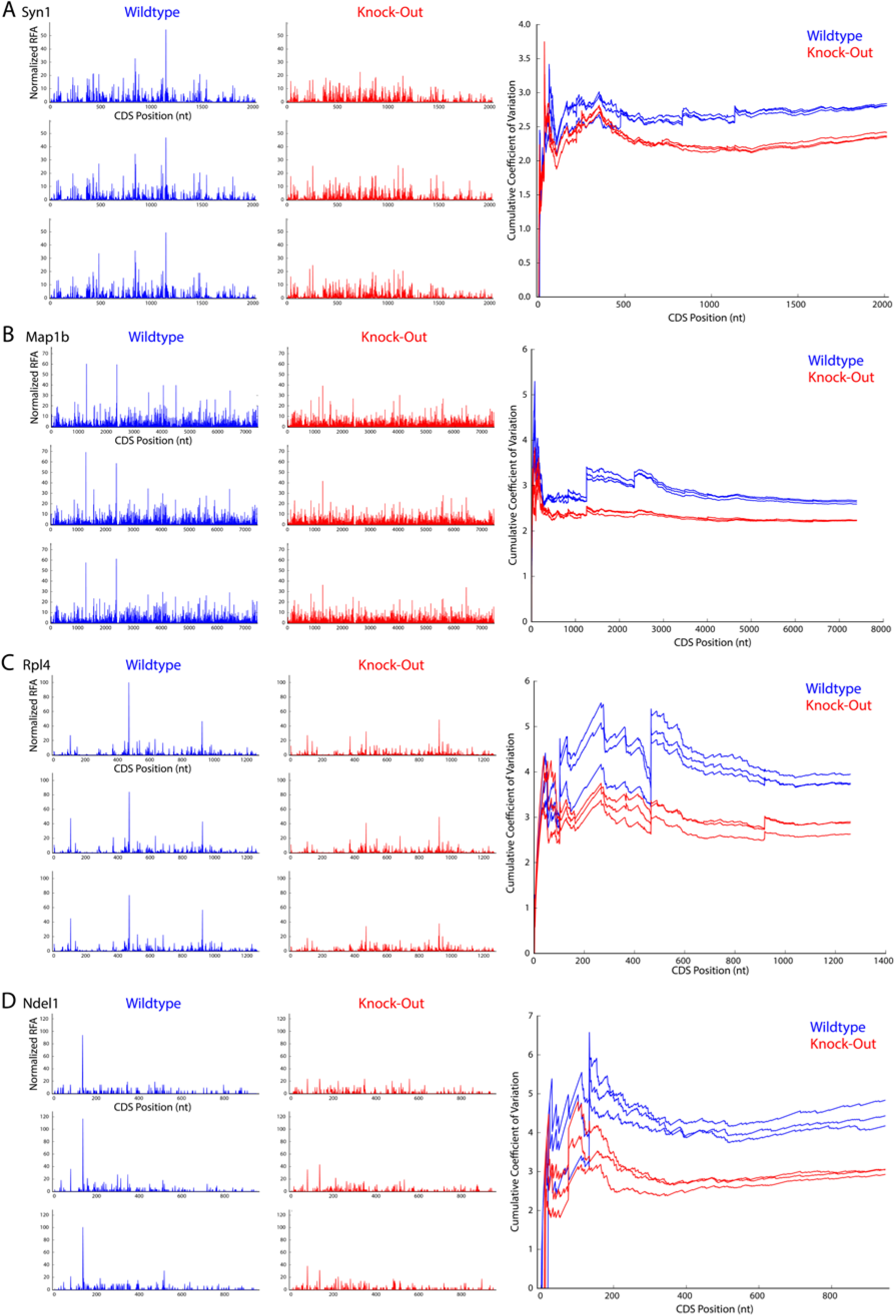
At left, nucleotide-resolution plots of P-site occupancy for genes *Syn1*, *Map1b*, *Rpl4*, and *Ndel1* (A-D, respectively) for three replicates of both wild-type (blue) and *Fmr1*-KO (red) mice. These plots are paired on the right with comparisons of the cumulative coefficient of variation (CV), calculated as the coefficient of variation for the coding sequence up to a given nucleotide position in the CDS. While *Syn1* and *Map1b* both exhibit decreased RFApm in *Fmr1*-KO mice, *Syn1* is a high-affinity binding partner of FMRP and *Map1b* is not; similarly, *Rpl4* and *Ndel1* both exhibit increased RFApm in Fmr1-KO mice but Rpl4 is a 5’-TOP gene and target of mTOR. For all these genes, the magnitudes of the “spikes” of reproducible, high-frequency P-site alignment, which represent pause sites, are significantly reduced in *Fmr1*-KO occupancy plots compared to their wild-type counterparts, and this reduction is reflected in a correspondingly diminished increase in cumulative CV at the pause site’s coordinate for *Fmr1*-KO replicates. The overall decrease in positional noise of aligned P-sites with FMRP loss, represented by the consistent gap in cumulative CV between genotypes at nearly all coordinates, is larger than that which can be explained by large pause-reductions alone.

## Discussion

Previous studies have shown that FMRP associates with polysomes and the protein-coding sequences of a large number of transcripts (Brown et al., 2001; Stefani et al., 2004). HITS-CLIP data indicate that FMRP has particularly high affinity for mRNAs involved in synaptic activity and appears to act as a translational brake, stalling ribosomes on these transcripts (Darnell et al., 2011). Prior work has also revealed interactions between FMRP and the translation initiation machinery (Napoli et al., 2008; Santini et al., 2017). Nonetheless, genome-wide measurements of protein synthesis with the resolution to analyze both translation elongation and RFApm have not been undertaken in the brains of *Fmr1*-KO mice.

We characterized the translational landscape in the cortex of *Fmr1*-KO mice at a crucial time in postnatal brain development. By P24, the mouse brain has reached its peak synaptic density and significant pruning of excitatory synapses is taking place, a process known to be dysregulated broadly in autism spectrum disorders (Tang et al., 2014) and specifically in FXS (Comery et al., 1997; He and Portera-Cailliau, 2013). Loss of FMRP-mediated regulation of protein synthesis may be critically linked to the synaptic plasticity and dendritic spine phenotypes observed in FXS (Darnell and Klann, 2013). We discovered a remarkably uniform trend in the RF abundance of FMRP’s high affinity binding partners with nearly all of the top 200 FMRP-bound transcripts showing a significant reduction in RFApm in *Fmr1*-KO mice (**Figure 2B**). This result is surprising because proteins encoded by many of these mRNAs have been shown to be more highly expressed in *Fmr1*-KO mice (Tang et al., 2015). Importantly, reduction in ribosome density was not a global effect. For example, the 5’TOP motif-containing mRNAs, which are comprised mainly of ribosomal protein- and translation factor-encoding transcripts, were enriched among genes with increased RFApm (**Figure 2C**). These genes are known to be controlled at the level of translation initiation by 4E-BP and eIF4E, the latter of which is sequestered by an FMRP-mediated complex.

Despite these clear patterns, RFApm is a complicated metric. In many studies, it is interpreted as a measure of translation efficiency that primarily reflects translation initiation. However, this interpretation assumes that initiation is rate-limiting and elongation rates are uniform (Arava et al., 2005). Given the potential role of FMRP in regulating translation elongation (Darnell et al., 2011), the apparent reduction in ribosome density for FMRP’s high-affinity binding partners (**Figure 2C**) may actually result from a relaxation of translational stalling in the absence of FMRP. We took advantage of the nucleotide resolution of ribosome profiling and characterized the noise in wild type and *Fmr1*-KO ribosome profiles with both codon motif- and gene-centric analyses. In both cases, we found a significant and global reduction in translational pausing in *Fmr1*-KO mice (**Figures 3-4**). As a genome-wide snapshot of translation in the cortex of *Fmr1*-KO mice *in vivo*, our results expand on previous *in vitro* measurements of ribosome stalling on select mRNAs using puromycin run-off (Darnell et al., 2011) and elongation rate using the ribosome transit time assay (Udagawa et al., 2013). We observed decreases in ribosomal pausing for the FMRP high-affinity binding partners, which exhibited a reduction in RFApm, and for the 5’TOP motif-containing mRNAs, which showed an increase in RFApm (**Supplementary Figure 2**). We note that our results do not formally rule out the possibility that the FMRP-associated mRNAs are also differentially regulated at the level of translation initiation. However, these results are consistent with a model in which FMRP loss dysregulates ribosomal pausing across a large number of transcripts, and that competition between initiation- (e.g., through FMRP-mediated sequestration of EIF4E) and elongation-level regulation results in disparate alterations in RFApm for certain genes.

We suggest that therapeutic strategies for FXS should carefully consider the consequences of globally altered protein synthesis. Recent evidence suggests that enhanced translation of certain mRNAs in *Fmr1*-KO mice may represent compensatory changes and that enhancing their function may ameliorate disease phenotypes (Thomson et al., 2017). Importantly, our study does not assess whether translational alterations in *Fmr1*-KO mice are caused by direct loss of FMRP function or by secondary effects arising due to continued absence of FMRP during neural development. A critical aspect is that neuronal activity may be tightly coupled to translational regulation. Several recent studies found translational repression of neuronal mRNAs following fear conditioning *in vivo* (Cho et al., 2015), and of FMRP binding partners following KCl depolarization *in vitro* (Dalal et al., 2017). Given extensive evidence of cortical hyperexcitability (Gibson et al., 2008; Hays et al., 2011) and dysregulation of GABAergic neurotransmission in *Fmr1*-KO mice (Paluszkiewicz et al., 2011), it is possible that the downregulation of RFApm we observed in FMRP binding partners (**Figure 2C**) is linked to increased cortical activity. We found enhanced translation of immediate early genes such as *Arc* and *Fos* as well as decreased translation of *Gabra2* (**Figure 1C),** consistent with previous reports of decreased GABA_A_ receptor expression and GABA dysfunction in FXS (Braat et al., 2015; D’Hulst et al., 2006). Future studies using knockdown or conditional knockout of *Fmr1* may be necessary to disentangle the primary effects of acute FMRP loss from secondary alterations in neuronal physiology. Nonetheless, our study shows that *Fmr1* loss leads to widespread alterations in mRNA translation, particularly at the level of elongation, during the developmental period of cortical synaptic refinement.

## Acknowledgements

We acknowledge the following core facilities for technical support: The JP Sulzberger Columbia Genome Center and the Institute for Comparative Medicine. PAS, GT, and D.S. were supported by grant 345915 from the Simons Foundation. PAS was supported by K01EB016071 from NIH/NIBIB. D.S. was supported by R01DA07418 from NIH/NIDA, R01MH108186 from NIH/NIMH, and the JPB Foundation. GT was supported by W81XWH-16-1-0263 from DOD and K01MH096956 from NIH/NIMH. NJH was supported by F31NS089106 from NIH/NINDS.

## Author Contributions

P.A.S., D.S., and G.T. conceived the project and designed the experiments. S.D.S., H.L., B.D.H., and G.T. conducted the experiments. S.D.S., J.B.M., B.D.H., N.H. and P.A.S. analyzed the data. All authors wrote and edited the manuscript.

## Data Availability

The ribosome profiling and RNA-Seq data have been deposited in the Gene Expression Omnibus (GEO) under accession GSE114064.

## Experimental Procedures

### Mice

*All mice were in* C57BL/6J background. *Camk2a*-cre-RiboTag mice were generated by crossing *Camk2a*-cre (JAX 005359) mice with RiboTag mice (JAX 011029) as reported previously (Hornstein et al., 2016). *Camk2a*-cre heterozygotes were crossed to RiboTag mice to obtain Rpl22^flox/flox^;*Camk2a*-cre^+/-^ mice, which were further crossed to Fmr1^−/y^ mice (Jax 00325) to generate Fmr1^X+/X-^;Rpl22^flox/flox^;*Camk2a*-cre^+/-^ females. The Fmr1^X+/X-;^Rpl22^flox/flox^;*Camk2a*-cre^+/-^ females were then bred to Rpl22^flox/flox^ males to obtain Fmr^1-/y^;Rpl22^flox/flox^;*Camk2a*-cre^+/-^ mice and Rpl22^flox/flox^;*Camk2a*-cre^+/-^ control littermates.. Throughout the manuscript, we refer to the Fmr1^−/y^;Rpl22^flox/flox^;*Camk2a*-cre^+/-^ mice as *Fmr1*-KO and the Rpl22^flox/flox^;*Camk2a*-cre^+/-^ mice as wild type. All experiments we conducted at postnatal day 24 (P24). All mouse experimental procedures were reviewed and approved by Columbia University Medical Center Institutional Animal Care and Use Committee.

The mice were genotyped with the following primers for Cre: GCG GTC TGG CAG TAA AAA CTA TC (transgene), GTG AAA CAG CAT TGC TGT CAC TT (transgene), CTA GGC CAC AGA ATT GAA AGA TCT (internal positive control forward), GTA GGT GGA AAT TCT AGC ATC ATC C (internal positive control reverse), and the following primers for RiboTag: GGG AGG CTT GCT GGA TAT G (forward), TTT CCA GAC ACA GGC TAA GTA CAC (reverse). The primers for Fmr1KO mice were: CAC GAG ACT AGT GAG ACG TG (mutant forward); TGT GAT AGA ATA TGC AGC ATG TGA (wild type forward); CTT CTG GCA CCT CCA GCT T (reverse)

### Tissue processing for RNA sequencing and Ribosome profiling

Brain tissue was processed as described previously (Hornstein et al., 2016). Briefly, snap-frozen frontal cortex (n=4 mice/genotype for RNA-Seq and n=3 mice/genotype for ribosome profiling, sample weight ~25mg) was disrupted using a Dounce homogenizer in 1mL of polysome lysis buffer (20 mM Tris-HCl pH 7.5, 250 mM NaCl, 15 mM MgCl_2_, 1mM DTT, 0.5% Triton X-100, 0.024 U/ml TurboDNase, 0.48 U/mL RNasin, and 0.1 mg/ml cycloheximide). Homogenates were clarified by centrifugation at 14,000 x g for 10 min at 4°C. Supernatant was collected and used for RNA-Seq and ligation-free ribosome profiling.

### RNA-Seq library construction

Total RNA was isolated from brain lysates using a Qiagen RNeasy kit (cat no. 74104) and ribosomal RNA was depleted using the Ribo-Zero rRNA removal kit from Illumina (Cat no. MRZH11124) according to the manufacturer’s instructions. rRNA depleted total RNA samples were converted to a strand-specific sequencing library using the NEBNext^®^ Ultra™ Directional RNA Library Prep Kit from Illumina (Cat no.E7420S). There were a total of four RNA-Seq libraries generated for each genotype, with each library originating from a different animal. RNA-Seq libraries were quantified using Qubit fluorometer (ThermoFisher) and library size was measured using an Agilent Bioanalyzer.

Sequencing of eight RNA-Seq libraries was performed on an Illumina NextSeq 500 desktop sequencer with a read length of 75 bases. Approximately 20 to 50 million demultiplexed, pass-filtered, single-end reads for each sample were obtained.

### Ligation-free ribosome profiling

Ligation-free ribosome profiling libraries were prepared from dephosphorylated foot-prints (~ 28-34 nucleotides in length) using a commercially available kit (SMARTer small RNA-Seq Library Preparation Kit, Clontech, Cat no. 635029) following manufacturer’s instructions (Hornstein et al., 2016). We performed library purification with AMPure XP beads (Beckman Coulter). Libraries were quantified using the Qubit dsDNA High-Sensitivity kit (Life Technologies) and library size was verified with the High-Sensitivity Bioanalyzer DNA chip (Agilent Technologies). Sequencing of six ribosome profiling libraries was done on an Illumina NextSeq 500 desktop sequencer with a read length of 50 bases. We obtained between 20 to 50 million demultiplexed, pass-filtered, single-end reads for each sample.

### High-Throughput Sequencing Data Processing

Bioinformatics analysis was performed following a protocol from Hornstein et al 2016 (Ingolia et al., 2012) with minor modifications. Ribosome profiling libraries were processed by removing the first 4 and last 10 positions of each sequenced read with the following command to fastx-trimmer:

fastx_trimmer -f 4 -l 40 -Q33 -i INFILE -o OUTLFILE
following which we trimmed remaining poly(A) sequence from the 3’ end, discarding trimmed reads shorter than 25 nucleotides. Libraries were then depleted of ribosomal RNA by alignment to an rRNA reference library comprised of rRNA sequences from mm9 with bowtie2, allowing for one alignment error. Unaligned reads were retained and aligned to the mm10 assembly of the mouse genome and Gencode-annotated transcriptome with STAR (Dobin et al., 2013). Alignments to the exons and coding sequences (CDS) of genes were counted with the featureCounts (Liao et al., 2014) program from the subread suite, yielding between 4 and 10 million reads uniquely mapped to the CDS per ribosome profiling library.

### Statistical Analysis of RNA Expression, Ribosome Footprint Abundance, and Ribosome Footprnt Abundance per mRNA (RFApm)

We used DESeq2 (Love et al., 2014) to analyze differential expression from uniquely aligned RNA-Seq reads and differential ribosome footprint abundance from ribosome profiling reads that aligned uniquely to the CDS of each gene. We used the generalized linear model in RiboDiff (Zhong et al., 2017) to analyze differential ribosome footprint abundance per mRNA (RFApm). For this analysis, only reads that aligned uniquely to the CDS were used for both RNA-Seq and ribosome profiling. We used the Java implementation of gene set enrichment analysis (GSEA) (Subramanian et al., 2005) to assess the statistical enrichment of gene ontologies. Specifically, we pre-ranked each gene by fold-change and used “classic” mode to compute normalized enrichment scores and corrected p-values for gene sets in the MSigDB C5 gene ontology collection.

### Codon Motif-Level Analysis of Pausing

Ribosome profiling libraries were first aligned to the transcriptome using the –quantmode TranscriptomeSAM option in STAR v2.5 as follows:

STAR --readFilesCommand zcat --genomeDir STAR_INDEX --runThreadN 12 --outSAMtype BAM SortedByCoordinate --readFilesIn INFILE --outSAMprimaryFlag AllBestScore --outSAMattrIHstart 0 --quantMode TranscriptomeSAM --outFileNamePrefix OUTFILE

Transcriptome-aligned libraries were then filtered by removing reverse-complemented (SAM flag 272 or 16), suboptimal, and non-CDS-aligned reads.

We chose one representative transcript and coding sequence for each gene by summing counts for all transcripts independently, then choosing the transcript with the highest sum of counts for each gene. To reduce reads from ~28-30nt footprints to A-site locations, we used the *psite* script from the plastid library for ribosome profiling analysis (Dunn and Weissman, 2016). This script calculates the location of a ribosomal P-site relative to the 5’ end of a footprint based on its length; increasing the calculated P-site offset by 3 nucleotides yields the A-site offset. We obtained codon occupancy profiles by summing over A-sites overlapping the 0, +1, and −1 nucleotide positions relative to the codon start, then merged them by summation across samples within either condition (wild-type or *Fmr1*-KO), collapsing six samples to two overall profiles with greatly increased coverage. We then limited the set of transcripts under consideration to those with mean coverage of at least 0.1 A-sites per codon for the first 150 codons in both profiles, yielding 8,967 total transcripts, and calculated pause scores for all but the first and last 10 codons within each.

Ribosome pause scores were calculated following the approach described by Woolstenhulme et al (Woolstenhulme et al., 2015), modified to correct for potential differences in splicing across profiles in line with Ishimura et al (Ishimura et al., 2014). We calculated context-specific pause scores for every codon of every coding sequence by dividing the codon’s ribosome occupancy by the maximum of three values: the mean occupancy of the first 150 codons of the transcript and the median occupancies of the five codons 5’ and 3’ to the codon in question. To obtain a mean pause score for each amino acid, we averaged scores across all occurrences of codons encoding that amino acid residue; di- and tri-amino acids with a minimum of 100 occurrences across the transcripts considered were similarly summarized. For mono-, di-, and tri-amino acid datasets, we performed a Mann-Whitney U-test to determine statistical significance of the difference in the distributions of pause scores between genotypes.

### Gene-Level Analysis of Translational Pausing

We used Ribo-TISH (Zhang et al., 2017) to determine the ribosome P-site offsets for each fragment length and P-site ribosome profiles for each transcript in our ribosome profiling data. For the initial quality control step, we used the following command:

ribotish quality -b BAMFILE -g GTF -p 16

followed by a prediction step with:

ribotish predict -b BAMFILE -g GTF -f GENOME_FASTA -o OUTPUT_FILE -p 16 –transprofile PROFILE_OUTPUT_FILE –framebest –seq –aseq

We then restricted our analysis to annotated ORFs, and for each isoform of each gene, we computed the mean coverage (number of ribosome footprints per codon) and the coefficient of variation (CV) in coverage (standard deviation in the number of ribosome footprints per codon divided by mean). For each gene, we selected the isoform with the lowest CV. Isoforms with extremely non-uniform coverage, which can result from low usage or exclusion of a subset of exons, are typically not the dominantly expressed isoform. Finally, as described under Results, we fit Equation 1 to a plot of log_2_(CV) vs. log_2_(mean coverage) to assess the genome-wide dependence of noise along the CDS on coverage using the *curve_fit* function in SciPy.

## Supplementary Information for “Widespread Alterations in Translation Elongation in the Brain of Juvenile *Fmr1* Knock-Out Mice”

**Supplementary Figure 1.**
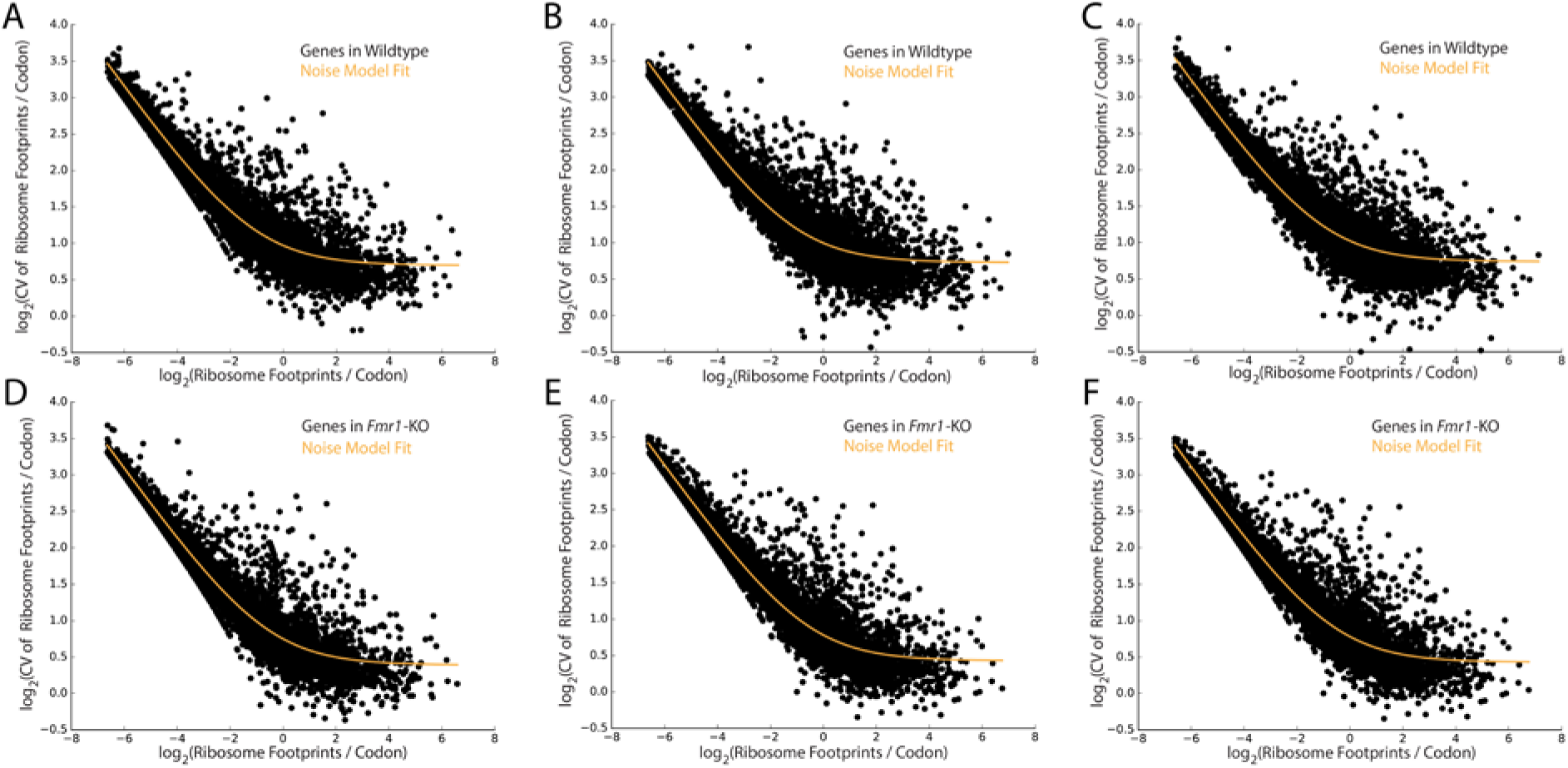
Coefficient of variation vs. mean of the number of ribosome footprints per codon across genes for A)-C) wildtype and D)-F) Fmr1-KO mice with fit to Equation 1.

**Supplementary Figure 2.**
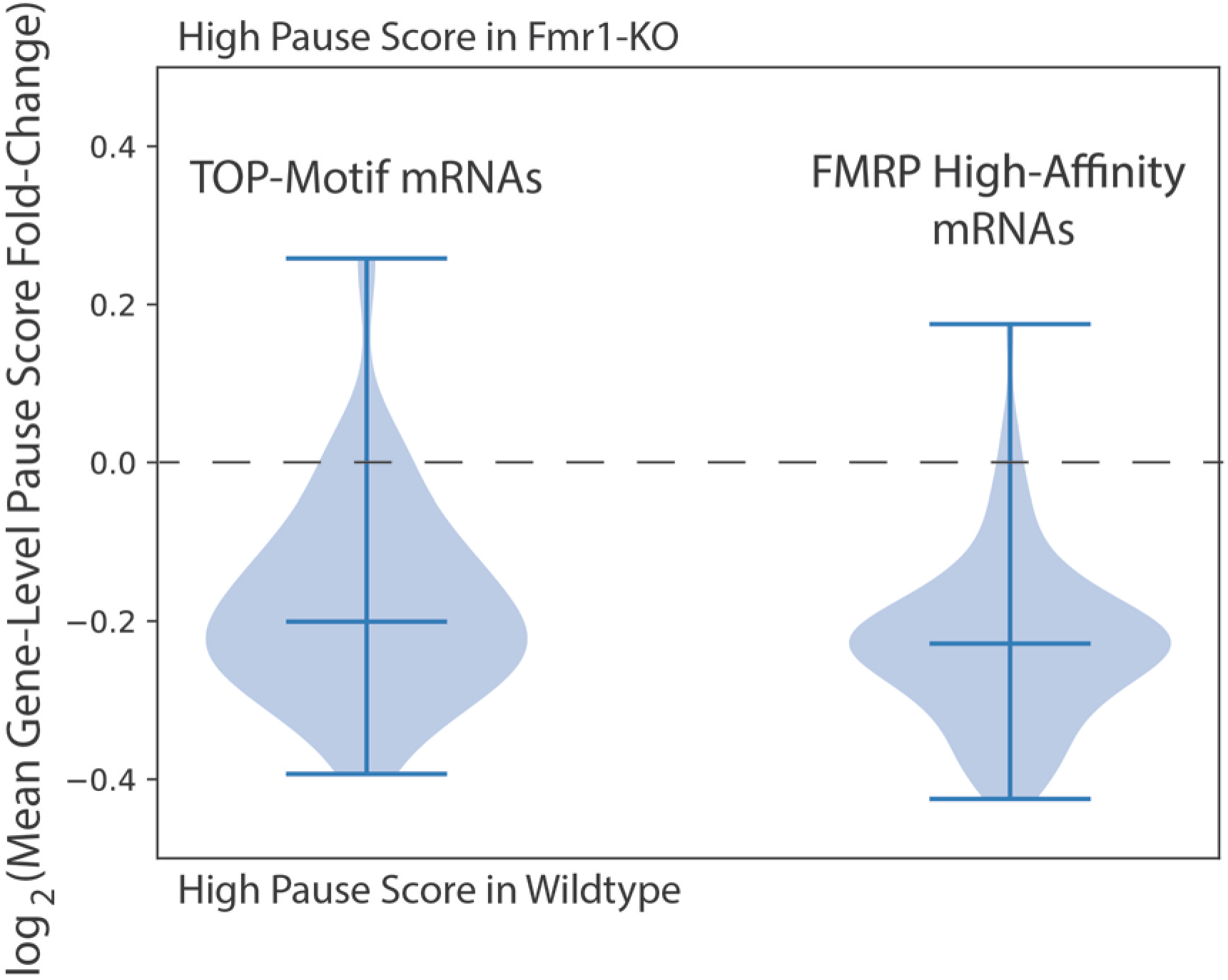
Distributions of fold-change in gene-level pause scores averaged across replicate experimental batches (each with a wildtype and Fmr1-KO mouse). The score is defined as the coefficient of variation divided by its expectation value from the noise model of the wildtype mouse (from the fit to Equation 1 shown in Supplementary Figure 1). We computed distributions for the TOP motif-containing mRNAs and the top 200 high-affinity FMRP binding partner mRNAs to show that nearly every gene in these two groups exhibits reduced pausing in Fmr1-KO mice.

**Supplementary Table 1.** Output of DESeq2 differential expression analysis comparing CDS-aligned ribosome footprint abundances between *Fmr1*-KO and wildtype mice.

**Supplementary Table 2.** Output of DESeq2 differential expression analysis comparing RNA-Seq profiles between *Frm1*-KO and wildtype mice.

**Supplementary Table 3.** Output of RiboDiff differential translation analysis comparing RFApm between *Fmr1*-KO and wildtype mice.

